# Chicken shank color determined by *Inhibition of dermal melanin* (*ID*) is mediated by a structural variation regulating *CDKN2A* expression

**DOI:** 10.1101/2024.12.24.630247

**Authors:** Jingyi Li, Lei Wang, Sendong Yang, Xin Zhou, Qinli Gou, Jinping Cai, Hongrui Yang, Qiaohua Wang, Shijun Li

## Abstract

It is well established that the color of a chicken’s shank is primarily determined by four genetic loci. Among these, the *Inhibition of dermal melanin* (*ID*) locus, which suppresses black pigmentation in the dermal layer of the shank, is the sole sex-linked mutation and its molecular mechanisms remained elusive. In this study, a resource family with segregating shank colors was constructed. A genome-wide association study utilizing FarmCPU software identified a top-associated SNP located on the Z chromosome. Subsequent linkage mapping further refined the candidate region, enabling the screening of the candidate structural variation. The candidate structural variation is associated with the yellow shank and characterized by a 143 bp deletion accompanied by a 2 bp insertion. Within the same Topologically Associating Domain, only the *CDKN2A* gene showed differential expression. Functional studies, including CRISPR/Cas9-edited cells, provided evidence that this mutation regulates *CDKN2A* transcription and is responsible for the *ID* shank color in chickens. The absence of melanocytes is likely due to their apoptosis. This study completes the puzzle of chicken shank color genetics and paves the way for the application of the *ID* mutation in the auto-sexing of chicks which is intensively needed in layer and broiler industries.

## Introduction

Shank color is one of the most conspicuous physical traits of domestic chickens, leading to a strong human preference for selecting chicken breeds with specific shank colors. For example, domestic chickens predominantly evolved from the red jungle fowl, yet the yellow shank mutation, originating from the gray jungle fowl, is now prevalent in most chicken breeds globally (Eriksson et al. 2008). Additionally, chickens with green or cyan shanks are favored by Chinese consumers, as they are perceived to signify high-quality chicken meat and eggs (Yang and Jiang 2005). Following these intensive breeding practices, the inheritance of chicken shank color has been extensively studied, resulting in the identification and establishment of four key genetic loci (Fig. 1).

**Figure 1.**
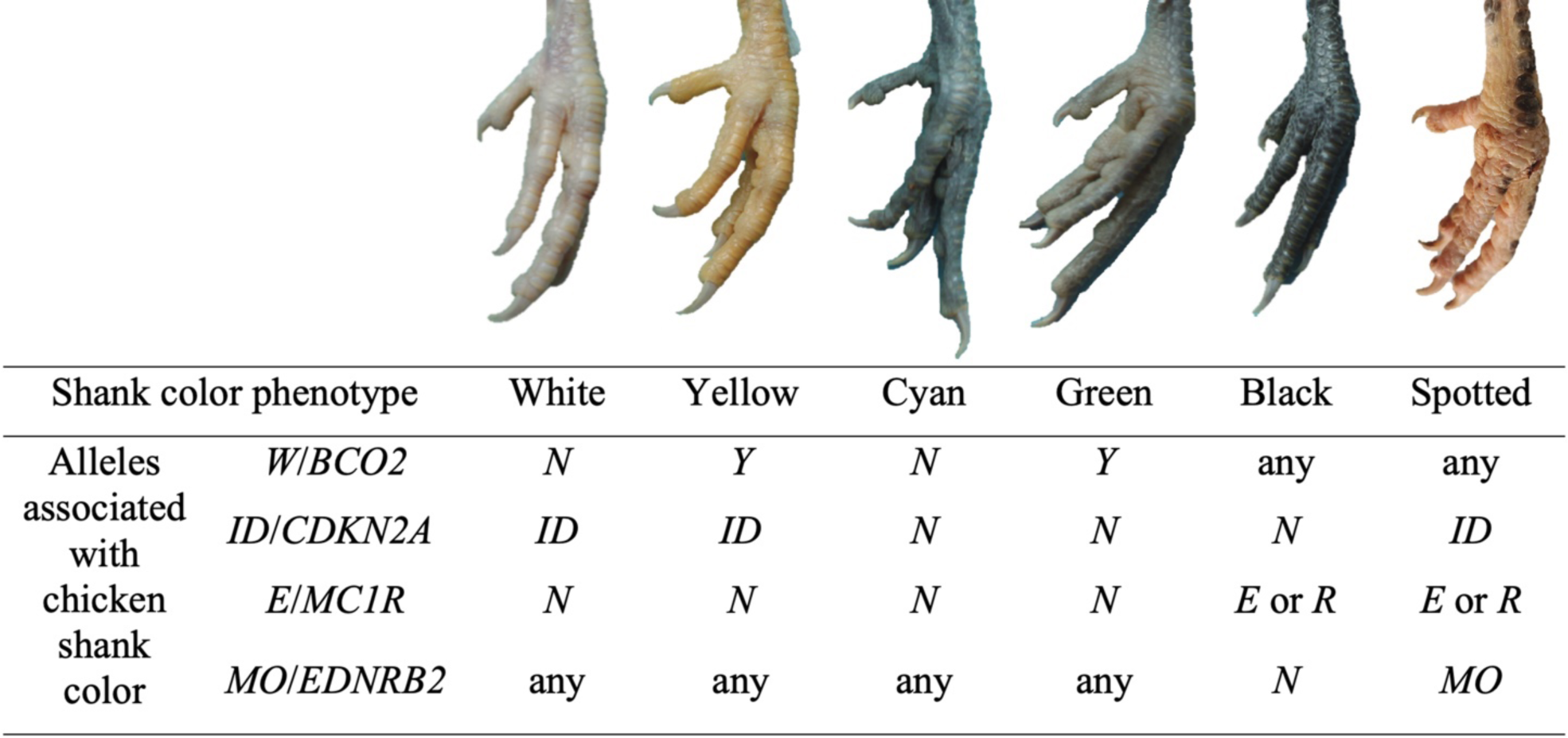
Variation in chicken shank color and associated genes. The photographs above were taken by Lei Wang. The table below is based on the review by Smyth (1990), with the addition of corresponding genes as identified in subsequent studies (Eriksson et al. 2008; Kerje et al. 2003; Kinoshita et al. 2014). *CDKN2A* was identified in the current study.

The *Yellow skin* (*W*) locus is associated with regulatory mutation(s) of the *BCO2* gene, located on chromosome 24 (Eriksson et al. 2008). Prior to the current study, the *Inhibition of dermal melanin* (*ID*) locus was mapped to the distal end of the Z chromosome, but the causal gene remained unidentified (Andersson et al. 2020). The *Extension* (*E*) locus is associated with missense mutations in the *MC1R* gene on chromosome 11 (Kerje et al. 2003). Furthermore, the *Mottling* (*MO*) locus has been mapped to a missense mutation in the *EDNRB2* gene on chromosome 4 (Kinoshita et al. 2014). In addition, the *Fibromelanosis* (*FM*) mutation, necessary for the expression of dark shank color in newly hatched chicks associated with the *ID*\**N* allele (Zhang et al. 2015; Yin et al. 2001; Kang et al. 2002), was found to be caused by duplication of the *EDN3* gene on chromosome 20 (Dorshorst et al. 2011). Among the mutations associated with shank color, *ID* stands out as the only one without a known genetic marker and is unique as the only sex-linked mutation.

Sexing 1-day-old chicks is essential for layer production, as only females are retained for egg production. This practice is also prevalent in broiler production, where males and females are raised separately due to their distinct growth patterns. Currently, auto-sexing has largely replaced vent sexing in the chicken industry. Early or late feathering is the most popular auto-sexing method because it does not require specific plumage backgrounds, making it suitable for both white and colored feather chickens. However, this method necessitates experienced identifiers for accuracy and is challenging for machine-based auto-detection. Overcoming these disadvantages, plumage color mutations can be utilized for auto-sexing but are not applicable to white layers or broilers, which are the world’s most popular chicken lines. Overall, shank color emerges as the optimal trait for auto-sexing, as it can be detected by machines and is applicable to chickens with white plumage. Moreover, early sexing of chick embryos through shank color is feasible and suitable for high-throughput automatic detection. Most importantly, euthanizing male layer embryos rather than 1-day-old chicks enhances animal welfare. Despite these advantages for auto-sexing, the absence of a genetic marker has hindered the widespread application of the *ID* locus in the chicken industry.

Following the identification of sex-linked inheritance in chicken shank color in 1911 (Bateson and Punnett 1911), and the naming of the *ID* locus in 1935 (Knox 1935), numerous studies have been conducted with the goal of mapping the *ID* locus and identifying its genetic markers. The *ID* locus has been mapped to a region between 78-79 Mb on the Z chromosome (GalGal6) based on six studies (Dorshorst et al. 2010; Li et al. 2014; Tian et al. 2014; Xu et al. 2017; Cha et al. 2023; Perini et al. 2024). The chicken breeds with *ID* mutations varied across these five studies, including Silkie, Tibetan, Youxima, Gushi, Ogye, and Italian local breeds. However, none of these studies were able to narrow down the candidate regions to specific candidate genes.

To conclusively identify the gene and causal mutation for *ID* in chickens, our team conducted the building of a resource family, conducting a Genome-Wide Association Study (GWAS), performing linkage mapping, developing diagnostic tests, and transcriptomic analyses. Furthermore, we undertook a series of functional studies, notably CRISPR/Cas9-mediated gene editing in cell culture experiments. Ultimately, we identified a candidate structural variation (SV) that results in the upregulation of the *CDKN2A* gene in the shank skin, thereby explaining the *ID* and lighter shank color phenotype.

## Results

### Inheritance and expression patterns of chicken shank color

As mentioned previously, *FM* and *ID* are two single-locus mutations mapped to chromosomes 20 and Z, respectively. By crossing a black-shanked male (*FM*N*/*N*, *ID*N*/*N*) with a yellow-shanked female (*FM*N*/*N*, *ID*ID*/W), we obtained offspring consisting of 8 males and 9 females, none of whom exhibited dark shank color at hatching. However, around the 14th day, the shank color of the females began to gradually darken, starting from the top of the shank and progressing to the bottom, becoming completely dark (including the shank, feet, and toes) by the 30^th^ day. These were classified as transformed shanks, with genotypes presumed to be *FM*N*, *ID*N*, aligning with previous reports (Zhang et al. 2015; Yin et al. 2001; Kang et al. 2002). The males, on the other hand, maintained yellow shanks throughout. This observation supports the notion that a single recessive gene on the Z chromosome is responsible for the black shank trait.

The second population was constructed for gene mapping purposes by crossing males with genotypes *FM*FM*/*N*, *ID*ID*/*N*, and females with genotypes *FM*N*/*N*, *ID*ID*/W. Among the 482 offspring, all 245 males exhibited yellow shanks. At week 9, the ratio of yellow shank to black shank females was 1:1 (116:121), which aligns with expectations. Among the 121 black shank females, 76 showed black shanks at hatching, while 45 were classified as transformed shanks. The observed ratio of black to transformed shank females did not match the expected 1:1 ratio (*P* = 0.005), suggesting that factors other than *FM* may also influence early pigmentation in black shank chicks.

### GWAS identifies top SNP associated with shank color

A GWAS conducted using FarmCPU software revealed a top SNP associated with shank color on the Z chromosome at position 78,952,074 bp (T to C transition, GalGal6 assembly, see Supplemental Results). For further analysis, 86 female offspring from the resource family, comprising 64 with black shanks and 22 with yellow shanks, were genotyped for this SNP. The results demonstrated a complete association between this SNP and shank color, suggesting that the causal mutation is likely this SNP or a mutation in its immediate vicinity.

### Transcriptomic analysis reveals cellular processes associated with shank pigmentation

To explore the mutations surrounding the top SNP, we designed 62 pairs of primers for Sanger sequencing of the approximately 50 kb region flanking the SNP. No coding mutations associated with shank color were identified, suggesting that changes in gene expression rather than gene function are responsible for the *ID*. Consequently, RNA-seq was employed using shank skin samples to identify candidate genes. According to the results, the five categories of differentially expressed genes (DEGs) (Supplemental Results, Supplemental Fig. S1) were plotted into a Venn diagram (Fig. 3A).

**Figure 2.**
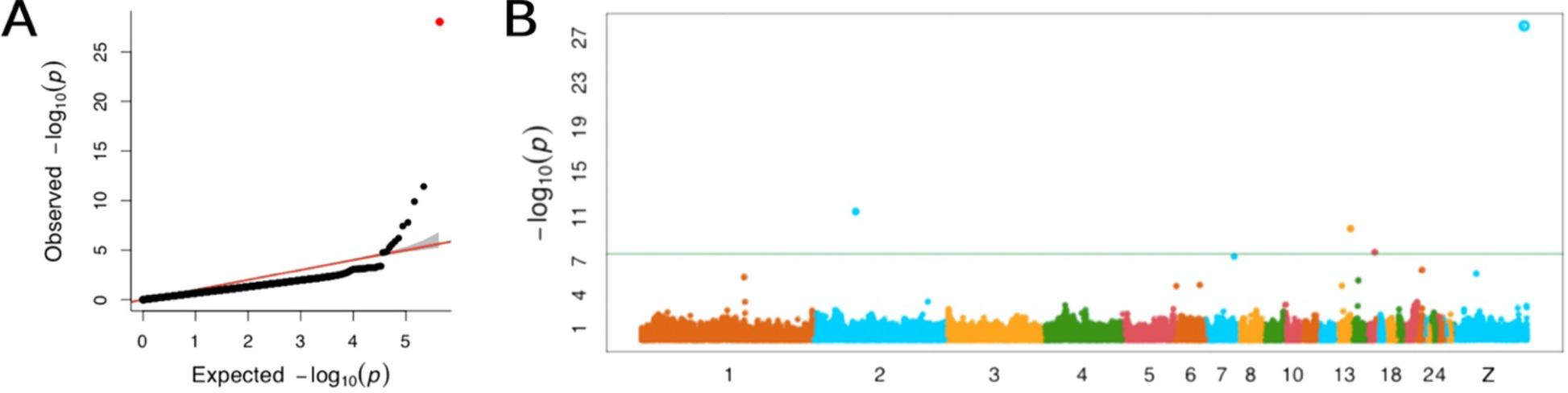
Results of GWAS for chicken shank color, conducted with FarmCPU. The GWAS for chicken shank color, executed using FarmCPU, included 76 cases and 286 controls. (*A*) The QQ plot from the GWAS is presented, with the strongest associated SNP indicated in red. (*B*) The genomic positions of the SNPs associated with chicken shank color, with references to the GalGal6 assembly.

**Figure 3.**
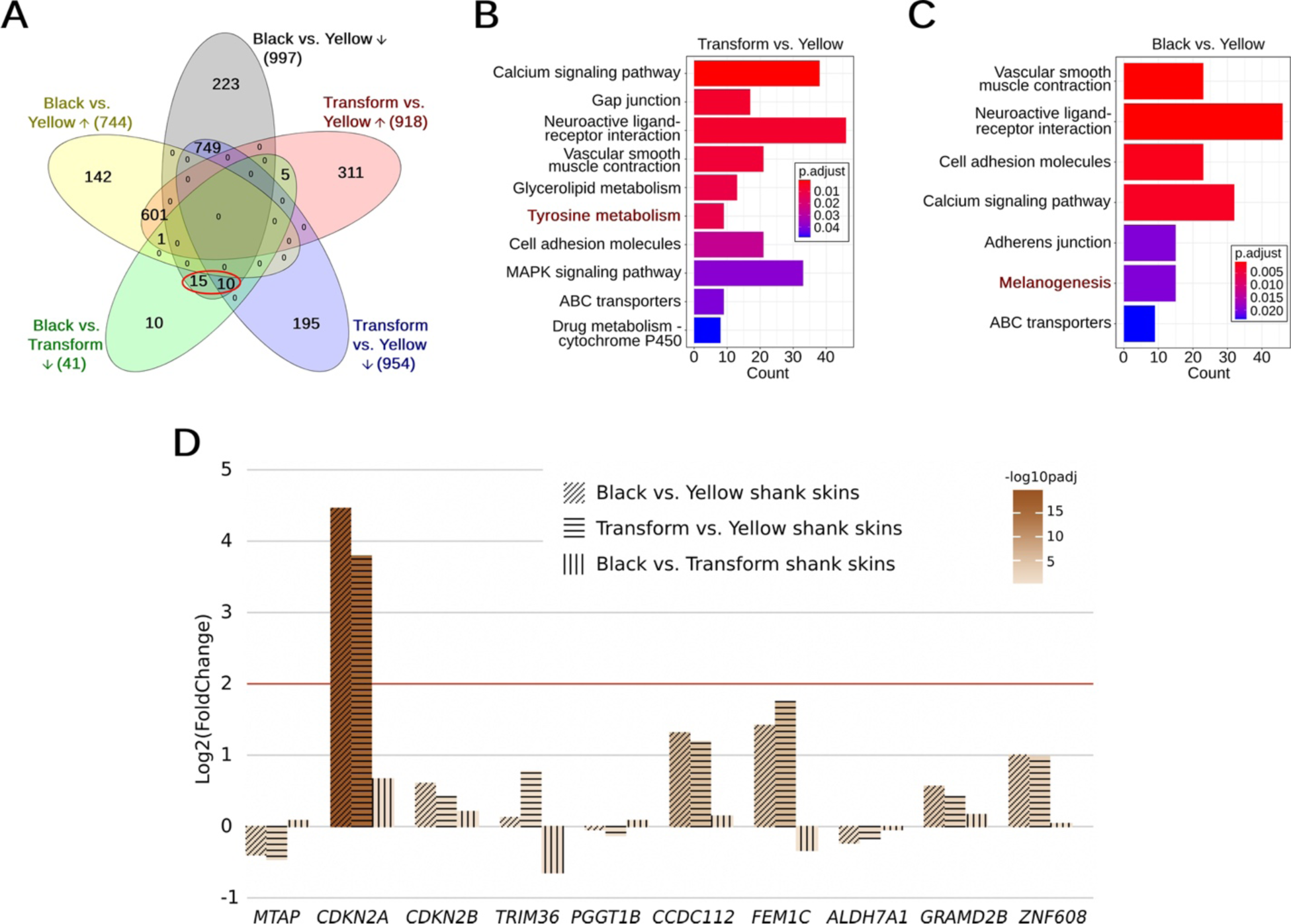
Transcriptomic analysis for chicken shank color. (*A*) A Venn diagram is presented for DEGs derived from various comparisons. Arrows pointing upwards and downwards indicate increased and decreased expressions, respectively. (*B*, *C*) KEGG pathway enrichment analysis is depicted, with pigmentation-related pathways highlighted in red. (*D*) *CDKN2A* is identified as the sole DEG among the 10 genes within the TAD, and it exhibits high expression in yellow shank skins. The arrangement of genes on the x-axis corresponds to their order on the Z chromosome.

The majority of DEGs were shared between the comparisons of black vs. yellow and transformed vs. yellow groups, with 601 upregulated and 749 downregulated DEGs. All non-shared DEGs (881 genes) had an average |log2foldchange| of 2.75, compared to 3.33 for the shared DEGs, indicating that a higher proportion of non-shared DEGs could be false positives due to lower fold changes. Additionally, the directionality of DEGs was consistent across the black and transformed groups, with no overlap between upregulated DEGs in black vs. yellow and downregulated DEGs in transformed vs. yellow comparisons, and vice versa. Collectively, these findings suggest that black and transformed samples exhibit very similar expression patterns.

To further investigate the expression patterns, DEGs from the three comparisons were mapped to Kyoto Encyclopedia of Genes and Genomes (KEGG) pathways. The tyrosine metabolism pathway was significantly enriched for DEGs in the comparison between transformed and yellow groups (Fig. 3B), primarily driven by three key genes essential for eumelanogenesis: *TYR*, *TYRP1*, and *DCT*. Concurrently, the melanogenesis pathway was enriched for DEGs between black and yellow groups (Fig. 3C), including the aforementioned three key genes and additional genes that regulate melanocytes or their pigmentation functions upstream, such as *WNT16*, *WNT3A*, *EDN1*, *EDNRB*, *EDNRB2*, *KITLG*, and *MC1R*.

A set of 25 genes was found to be exclusively highly expressed in the black group, that are the overlap between downregulated DEGs in both black vs. yellow and black vs. transformed comparisons (encircled in red in Fig. 3A). These genes encompass nearly all known genes responsible for melanin production, packaging, and transportation, including *MLANA*, *MC1R*, *TYR*, *TYRP1*, *DCT*, *PMEL*, *SLC45A2*, and *MLPH*. These pigmentation genes or melanocyte markers were scarcely expressed in the yellow group. In conclusion, our transcriptomic data indicates that yellow shank skins lack melanocytes, whereas melanocytes are present in both black and transformed shanks, but pigment production occurs only in black shank skins.

### The ID mutation is associated with the ectopic expression of CDKN2A in the shank

By integrating genomic and transcriptomic data, we investigated the expression patterns of genes surrounding the top SNP. Promoters or enhancers can regulate genes at a distance but are typically confined within the same Topologically Associating Domain (TAD). Due to chromosome folding, genes and elements within the same TAD can interact with each other but seldom with those in different TAD (Szabo et al. 2019).. According to publicly available FR-AgENCODE data (Foissac et al. 2019), the top SNP is located within a TAD (chrZ:78,739,756-80,339,755, based on liver and T cells) that contains 10 annotated genes: *MTAP*, *CDKN2A*, *CDKN2B*, *TRIM36*, *PGGT1B*, *CCDC112*, *FEM1C*, *ALDH7A1*, *GRAMD2B*, and *ZNF608*. Among these, *CDKN2A* was the only DEG across all three transcriptomic comparisons (with log2FoldChange values of 4.45 or 3.78 and adjusted *P*-values of 1.46 × 10^-20^ or 4.98 × 10^-18^ in comparisons between black and yellow groups or transformed and yellow groups, respectively, as shown in Fig. 3D), and it is exclusively expressed in yellow shank skins. Therefore, the ectopic expression of *CDKN2A* in yellow shank skin is likely responsible for the *ID* mutation.

### Identification of the candidate SV associated with ID

We then concentrated on the upstream region of *CDKN2A*, specifically the most proximal 10 kb within the previously Sanger-sequenced 50 kb region. Our analysis identified 5 top-associated SNPs in this area, but no associated SV were detected. Genotyping these 5 SNPs and conducting linkage mapping with 135 female offspring from the resource family revealed genetic distances between *ID* and each SNP of 1.46 cM, 0.73 cM, 0.73 cM, 0.73 cM, and 0 cM, respectively (from closest to *CDKN2A* to farthest; details of these SNPs are provided in Supplemental Table S1). This suggests that the causal mutation for *ID* may be located further from *CDKN2A* (see Supplemental Fig. S2 for a visual representation). Based on the recombination rate of 8 cM/Mb or higher at the end of the Z chromosome (Groenen et al. 2009), the causal mutation is likely to be positioned downstream but within approximately 87.5 kb of the 4th SNP, which is located at chrZ:78,923,391-79,010,891 (Supplemental Fig. S2). This linkage mapping outcome was corroborated by association tests and confirmed using a more advanced genome assembly (further details in the Supplemental Results).

Within the overlap of the linkage mapping region and the 50 kb Sanger-sequenced region, we identified two indels in an intergenic area, downstream of *TRIM36* and upstream of *CDKN2A*/*B*. The first indel is a 143 bp deletion (chrZ:78,942,980-78,943,123), which is replaced by two adenines (AA) and is strongly associated with yellow shanks. The same 86 females used for genotyping the top SNP were employed to genotype this candidate SV individually, and a complete association was found. The second indel was ruled out by diagnostic tests (Supplemental Results).

In the entire mapping population, consisting of 482 birds, and a breed panel comprising six breeds (48 individuals per breed, totaling 288 birds), the 143 bp deletion with a 2 bp insertion (the candidate SV) was genotyped individually. This SV was significantly associated with shank color in the mapping population (*P* = 3.2e-95, Fig. 4A), with a stronger association than the five SNPs used for linkage mapping (Supplemental Results). In the breed panel, complete associations were observed in White Leghorn, Houdan, Chahua, and Jingyang breeds. However, 6.3% of Yunyang Black-bone chickens (with black shanks) carried the mutation, and 18.6% of Jinghong No.1 layers (with yellow shanks) lacked the mutation. Nonetheless, the breed panel also demonstrated a significant association (*P* = 1.2e-54, Fig. 4B).

**Figure 4.**
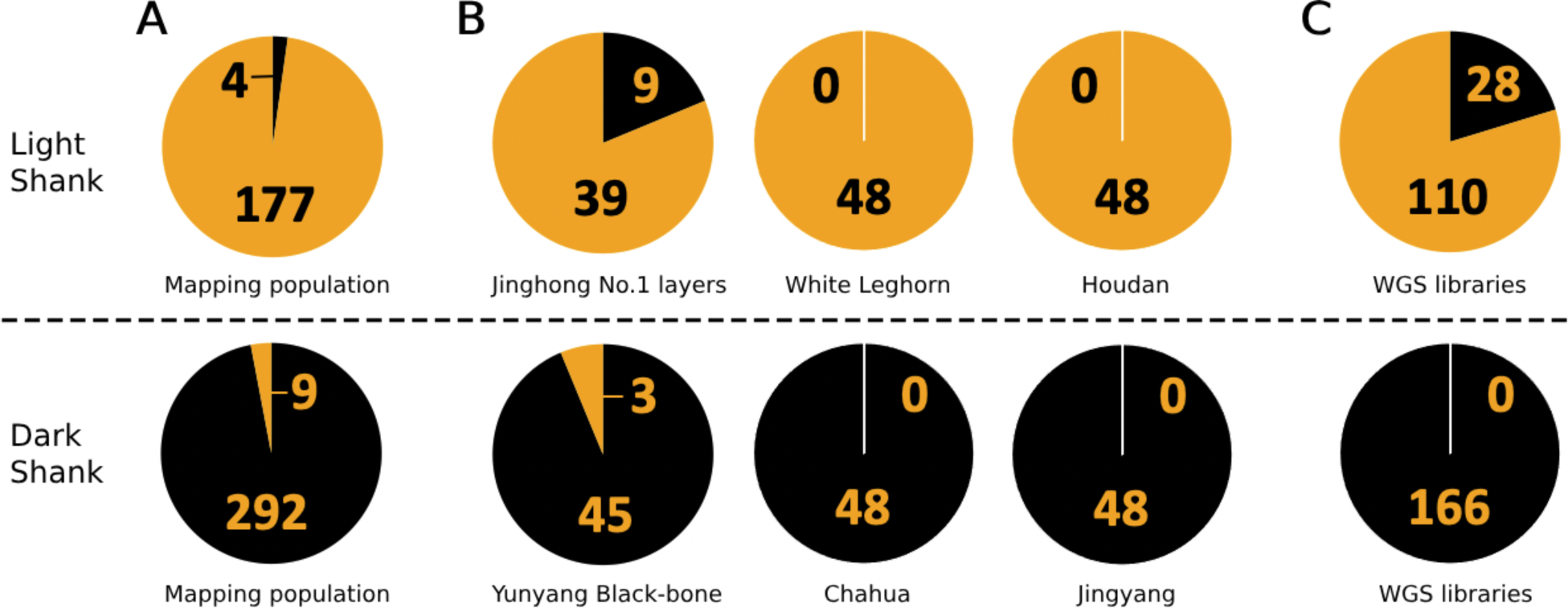
Diagnostic test results for the candidate SV in chickens with different shank colors. The numbers indicate the count of individuals for each genotype. Black segments of the pie charts represent the homozygous wild-type of the SV, whereas yellow segments denote the presence of at least one mutant allele copy. (*A*) Displays the results from the mapping population established in the current study. (*B*) Shows the findings from the breed panel. (*C*) Illustrates the results derived from WGS data.

The presence of the candidate SV was also examined in whole-genome sequencing (WGS) data, both from this study and from publicly available databases, totaling 304 individuals. These WGS data were obtained from chicken breeds with fixed shank colors. The results showed that the mutant allele was not detected in any of the 166 individuals from 14 breeds with dark shank colors, whereas it was found in 110 out of 138 individuals from 12 breeds with light shank colors, giving an allelic frequency of 80% (Fig. 4C, see Supplemental Table S2 and S3 for details).

Within a 20 kb region upstream and downstream of the candidate SV, 2,464 SNPs and SVs were identified from the 304 individuals. Apart from the candidate SV, 545 of these mutations were not present in the 166 individuals with dark shank colors, and their maximum allelic frequency in the 138 individuals with yellow shanks was only 5%. Consequently, the candidate SV remains the top candidate within the surrounding region.

### The candidate SV exhibits transcriptional regulatory effects

As the candidate SV is located upstream of *CDKN2A*, publicly available databases were utilized to predict its regulatory functions (Supplemental Results). To confirm the regulatory effects of the candidate SV, dual-luciferase reporter assays were conducted. The promoter activities of both the reference and mutant sequences were significantly higher than that of the mock control, with the mutant sequence exhibiting significantly higher promoter activity than the reference sequence (Fig. 5A). Similarly, the enhancer activities of both sequences were significantly higher than that of the mock control, although the mutant sequence displayed lower enhancer activity compared to the reference sequence (Fig. 5B).

**Figure 5.**
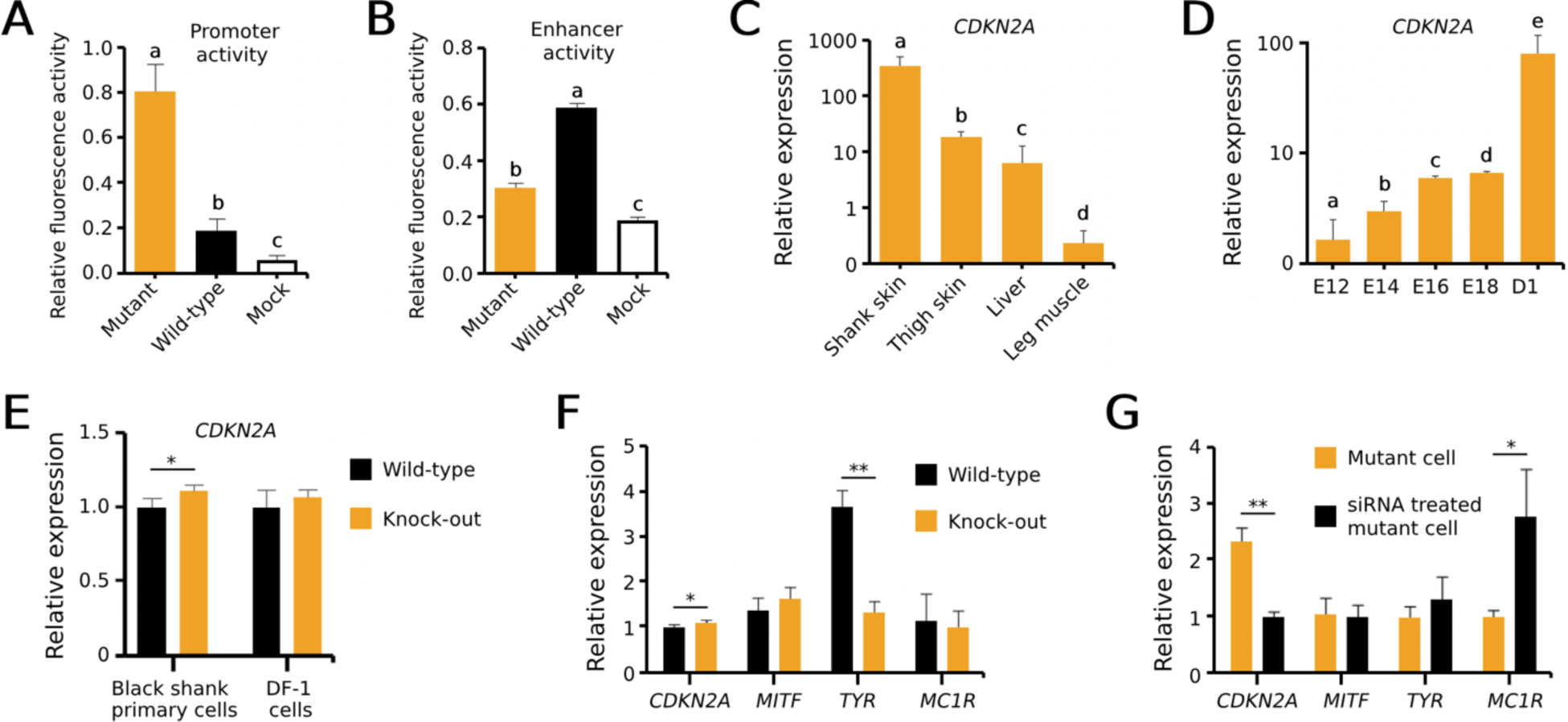
Regulatory effects of the candidate SV on *CDKN2A* expression. Data are presented as mean ± SEM. Distinct letters denote significant differences at the *P* < 0.05 level. An asterisk signifies a significant difference at *P* < 0.05, while double asterisks indicate highly significant differences at *P* < 0.01. (*A*, *B*) Dual-luciferase reporter assays disclose the transcriptional activities of recombinant plasmids representing distinct genotypes of the candidate SV (n = 3). (*C*) *CDKN2A* expression levels in various tissues of yellow shank female chicks at hatching (n = 3). (*D*) *CDKN2A* expression in shank skins of yellow shank female embryos across different developmental stages (n = 4). (*E*) Impact of knocking out the candidate sequence on *CDKN2A* expression in two different cell types (n = 3). (*F*) Impact of knocking out the candidate sequence on pigmentation-related gene expression in black shank skin cells (wild-type) (n = 3). (*G*) Impact of inhibiting *CDKN2A* expression on yellow shank skin cells (mutant) (n = 3).

### CDKN2A expression pattern in yellow shank skin during embryonic development

We further examined the expression pattern of *CDKN2A* using qPCR in embryos and chicks with yellow shanks. The results revealed that *CDKN2A* is expressed 3,422 times higher in shank skin than in muscle and at even higher levels compared to thigh skin and liver (Fig. 5C). Additionally, the expression of *CDKN2A* in shank skin increased as the embryo developed over time (Fig. 5D), aligning with the developmental pattern of melanoblasts in black shank embryos.

### In vivo infection not observed

To assess the feasibility of *in vivo* knockout of the 143 bp sequence, three different serotypes of adeno-associated virus (AAV) carrying enhanced green fluorescent protein (EGFP) were injected into chicks. However, no GFP fluorescence was observed in the tissue sections. Subsequently, these sections were subjected to immunohistochemistry, but no differences were observed between the injected and control groups (Supplemental Fig. S3). This suggests that the AAVs had low infection efficiency in chick dermal cells. Consequently, we proceeded to knock out the 143 bp sequence in cultured cells rather than pursuing *in vivo* knockout.

### Successful construction of double knock-out cells

A CRISPR/Cas9 double knock-out system was designed to target the deletion of the 143 bp sequence in wild-type cells. Following the synthesis of sgRNA, digestion, ligation, transfection, amplification, and extraction of plasmids, as well as sequencing (Supplemental Fig. S4), four types of knock-out vectors were successfully constructed. Upon transfection into DF-1 cells, all single and co-transfections displayed clear knock-out bands (Supplemental Fig. S5). We selected sgRNA3.2 and sgRNA5.2 for their highest mutation rate and proceeded with TA-cloning and sequencing. Among the 20 sequenced single-cell colonies, one showed a large fragment deletion in the candidate region, similar to the 143 bp deletion (Supplemental Fig. S6). Thus, the estimated mutation rate is 5%.

### Knock-out cells display altered CDKN2A and TYR expression

Primary shank dermal cells were isolated from six embryos with black shanks, and both these cells and DF-1 cells were genotyped for the candidate SV. The results indicated that the black shank cells were wild-type, while the DF-1 cells exhibited chimerism, with the majority being wild-type and a small portion carrying mutant alleles (Supplemental Fig. S7). This variability may be due to the non-clonal origin of our DF-1 cells. Using these two cell lines, we constructed double knock-out cells to artificially induce the candidate mutation in wild-type cells. In black shank cells, the knock-out significantly increased *CDKN2A* expression (*P* < 0.05), an effect not observed in DF-1 cells (Fig. 5E). Additionally, we analyzed the expression of melanocyte-specific genes, *MITF*, *TYR*, and *MC1R*, in black shank cells. No significant differences were found for *MITF* or *MC1R*; however, *TYR* expression was significantly reduced in knock-out cells (*P* < 0.01, Fig. 5F).

### Inhibition of CDKN2A leads to increased MC1R expression

Primary shank dermal cells from embryos with yellow shanks were collected and confirmed to be homozygous mutants for the candidate SV. Upon transfection with *CDKN2A*-targeting siRNA, *CDKN2A* expression was significantly inhibited (*P* < 0.01), resulting in an upregulation of *MC1R* (P < 0.05), while *MITF* and *TYR* expressions remained unchanged (Fig. 5G).

### Overexpression of CDKN2A induces apoptotic cell death

We successfully developed cell models with overexpression of *CDKN2A*, which was confirmed by qPCR (Fig. 6A). To assess the extent of apoptosis, Annexin V/PI flow cytometry was conducted (Fig. 6B). The results revealed that the apoptosis rate was significantly higher in cells overexpressing *CDKN2A* (Fig. 6C).

**Figure 6.**
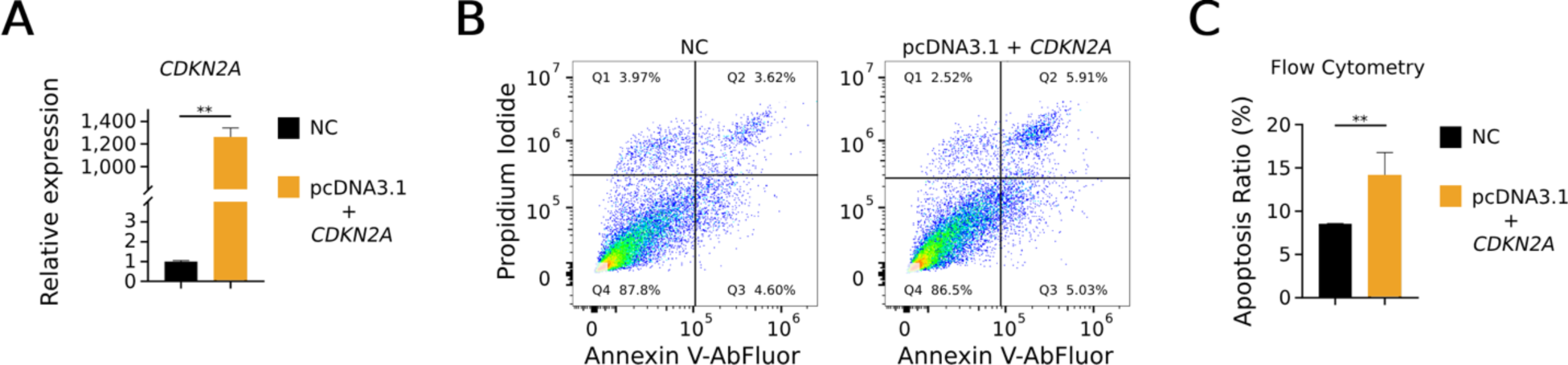
Apoptotic effects induced by *CDKN2A* overexpression. Data are presented as mean ± SEM. Double asterisks denote extremely significant differences at the *P* < 0.01 level. (*A*) *CDKN2A* expression levels were significantly higher in cells transfected with the overexpression vector (pcDNA3.1 + CDKN2A) compared to those with the negative control (NC) vector (pcDNA3.1), confirming the successful establishment of a *CDKN2A* overexpressing cell model (n = 3). (*B*) The proportion of apoptotic cells was assessed 48 hours post-transfection with either the *CDKN2A* overexpression vector or the NC vector, as measured by flow cytometry. (*C*) An increased rate of apoptosis was observed in cells overexpressing *CDKN2A*, suggesting that *CDKN2A* induces apoptotic cell death (n = 3).

## Discussion

### Establishing causality between the candidate SV and ID

To pinpoint the causal mutation for *ID*, a resource family was established in this study, followed by the application of FarmCPU for a GWAS. As anticipated, a single SNP emerged on the Z chromosome, rather than a peak with numerous markers. This characteristic of FarmCPU helped to exclude other weakly linked markers, but it also complicated the definition of the candidate region. Fortunately, this SNP is located within a known TAD, which thus served as the candidate region. Subsequently, transcriptomic data revealed that *CDKN2A* is the sole DEG in this region. After linkage mapping and mutation screening, a candidate SV, a 143 bp deletion with a 2 bp insertion, potentially regulating *CDKN2A* was identified. Its associations were confirmed in our resource family, breed panel, and WGS data. Additionally, flanking mutations were excluded as candidates based on the WGS data.

The location of this SV at chrZ:78,942,980-78,943,123 (GalGal6 assembly, same hereinafter) aligns with previously reported candidate regions for *ID*, with one exception. In 2010, two SNPs were reported to be significantly associated with ID: rs14779590 at chrZ:69,048,452 and rs14686603 at chrZ:79,154,166 (Dorshorst et al. 2010). Subsequently, Li et al. (2014) and Tian et al. (2014) reported three SNPs (rs315631308 at chrZ:78,914,305, rs317250881 at chrZ:79,556,958, and rs315850000 at chrZ:79,559,850) and two SNPs (rs16127903 at chrZ:73,599,275 and rs14685542 at chrZ:80,136,097), respectively. Xu et al. (2017) reported eight SNPs, spanning between rs16683912 at chrZ:79,418,742 and rs14685747 at chrZ:79,834,996. Cha et al. (2023) reported 13 SNPs between rs14686141 at chrZ:78,748,287 and rs14686097 at chrZ:79,326,198. Recently, Perini et al. (2024) reported multiple regions on chromosome Z associated with chicken shank color, including one region at chrZ:78,846,780-79,213,873 that contains *CDKN2A*. Some of these candidate regions do not overlap, potentially due to the lower quality of previous reference genome assemblies, particularly in the distal end of the Z chromosome, which is a highly repetitive region. Utilizing the GalGal6 genome assembly and *de novo* assemblies through advanced techniques (Zhu et al. 2023) along with intensive screening for variances, we have successfully identified the top candidate mutation for *ID*.

While we observed some exceptions in the association between this SV and *ID*, it is important to note that a portion of these discrepancies could be attributed to errors in phenotyping and genotyping. However, we noted that the highest ratio of unmatched samples originated from light shank populations, both in the breed panel and the WGS dataset. It has been reported that the *Dominant White* (*I*I*), *Sex-linked Barring* (*B*B*), and *FM*N* alleles can inhibit the expression of shank pigmentation (Knox 1935; Dorshorst and Ashwell 2009; Zhang et al. 2015), leading to light shank color in chickens that may still carry the *ID*N* allele. With the causal mutations of all these genes now identified, their epistatic interactions can be tested. Armed with this knowledge, it is possible to establish chicken lines that stably express different shank colors at the embryonic stage, enabling high-throughput automatic detection for sexing embryos prior to hatching.

### Molecular pathways in the formation of light shanks in chickens

We propose that *CDKN2A* is regulated by the candidate SV located 86 kb upstream. Our dual-luciferase reporter assay demonstrated that this SV possesses promoter activity, consistent with the ectopic expression of *CDKN2A* observed in yellow shank skin. However, we also noted a weaker enhancer effect for this SV compared to the wild-type. This contradiction could be attributed to two factors: 1. The SV, being a 143 bp deletion with a 2 bp insertion, introduces changes in more motifs (Supplemental Table S4) than SNPs or smaller SVs. Consequently, the increased promoter activity and decreased enhancer activity might be due to different motifs involved within the same SV. 2. Conflicting results have been reported using two different dual-luciferase reporter systems, cautioning against overreliance on these results (Wu et al. 2015). Despite these considerations, our double knock-out cells showed a significant increase in *CDKN2A* expression, providing more direct evidence of the regulatory effect of this SV.

The 86 kb distance is typically too great for a promoter element to directly interact with its target gene. Instead of a direct promoter-to-transcription start site (TSS) regulation, long-range promoter-promoter interactions are a more plausible mechanism in this context. The best predicted direct promoter for chicken *CDKN2A* is located 836 bp upstream of its TSS (Lizio et al. 2017). Sanyal et al. (2012) demonstrated in human cells that the peak frequency of promoter-promoter interactions occurs approximately 120 kb upstream of the TSS, with similar frequencies observed within the 0 to 200 kb range upstream of the TSS. Chromatin loops are likely involved in these long-range interactions. Makino and Fukaya (2024) reviewed three examples of long-range promoter-promoter loops in Drosophila, with distances ranging from 235 kb to approximately 6 Mb. Based on the distinct Hi-C signals (Supplemental Fig. S8), the distance between the 143 bp sequence and *CDKN2A* after chromatin folding is likely much less than 86 kb, permitting the 143 bp sequence to regulate *CDKN2A* expression (Fig. 7). This finding aligns with our observations in knock-out cells, where the candidate sequence was targeted for knockout using CRISPR/Cas9, leading to upregulation of *CDKN2A* in shank skin cells (Fig. 5G). The involvement of long-range interactions and potential differences in these interactions among different *ID* genotypes can be further examined using Hi-C.

**Figure 7.**
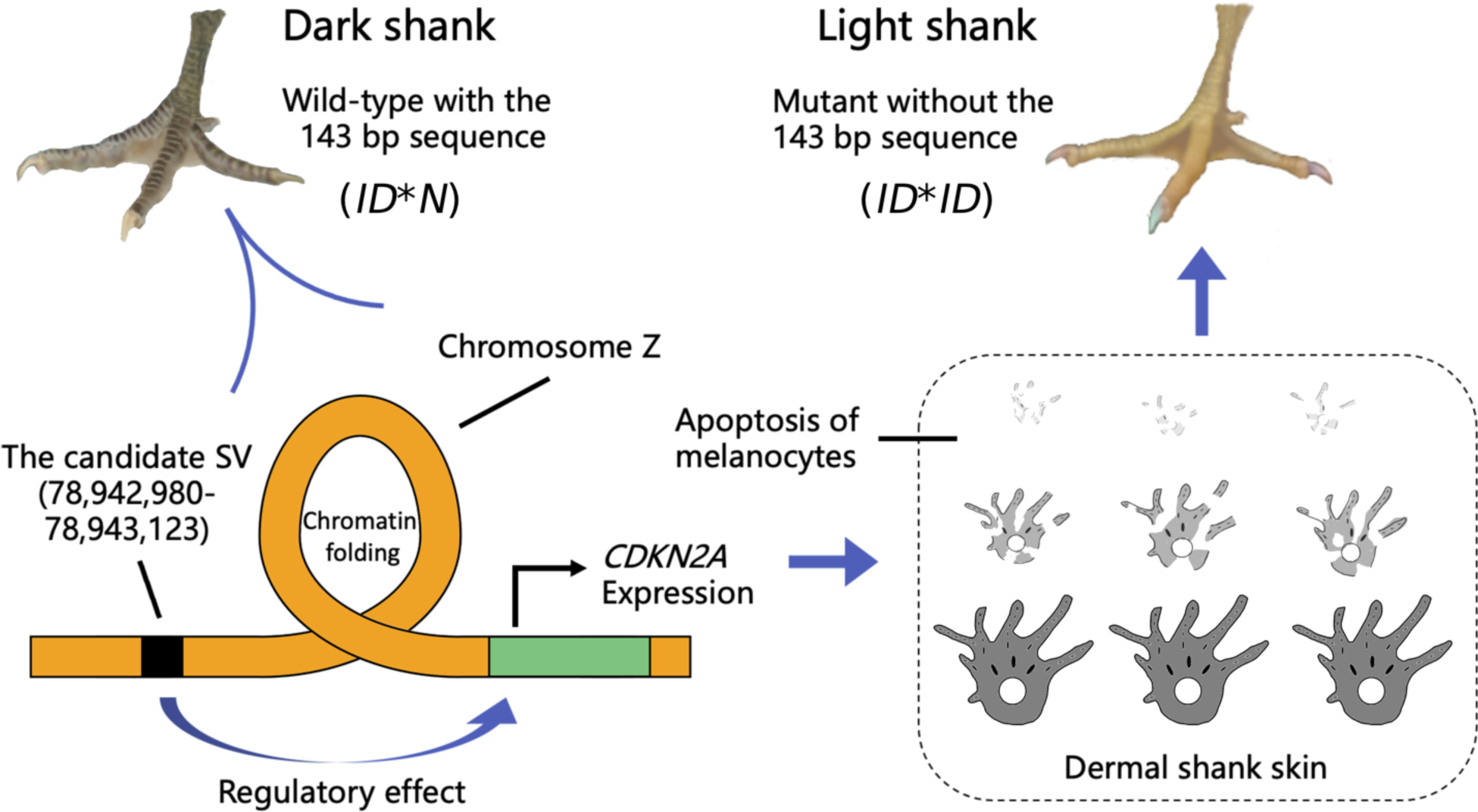
Schematic representation of the of the mechanism behind light shank color (*ID*\**ID*) in chicken. The 143 bp candidate sequence, found in chickens with dark shanks, is located 86 kb upstream of the *CDKN2A* gene. However, through chromatin folding, this candidate SV, a 143 bp deletion accompanied by a 2 bp insertion, becomes adjacent to *CDKN2A*, thereby enhancing gene expression. The ectopic expression of *CDKN2A* triggers apoptosis in melanocytes and their precursors, resulting in the absence of melanin in the light shank skin.

The upregulation of *CDKN2A* leads to apoptosis, which was supported by our cell overexpression experiment (Fig. 6) and also literatures (Ji et al. 2020; Chakraborty et al. 2021; Luan et al. 2021). We also showed the spatial-temporal expression pattern of *CDKN2A* in chicken embryos, indicating its role in shank pigmentation. However, variants of *CDKN2A* were not found to be associated with skin color in human (Ibarrola-Villava et al. 2010), while change of skin pigmentation was not detected in *CDKN2A* knockout mice according to International Mouse Phenotyping Consortium (IMPC) (Dickinson et al. 2016). This might suggest that the roles of CDKN2A in melanocytes are different in mammals and avians.

The upregulation of *CDKN2A* leading to apoptosis is supported by our cell overexpression experiments (Fig. 6) as well as by literature (Ji et al. 2020; Chakraborty et al. 2021; Luan et al. 2021). Our study also demonstrated the spatial-temporal expression pattern of *CDKN2A* in chicken embryos, highlighting its role in shank pigmentation. However, variants of *CDKN2A* were not found to be associated with skin color in (Ibarrola-Villava et al. 2010), and changes in skin pigmentation were not observed in *CDKN2A* knockout mice, according to the International Mouse Phenotyping Consortium (IMPC) (Dickinson et al. 2016). This could suggest that the roles of CDKN2A in melanocytes differ between mammals and avians.

Our RNA-seq data supports the hypothesis that the absence of melanin in yellow shank skin is due to apoptosis. Firstly, Caspase-3, a marker for apoptosis, was significantly more highly expressed in yellow shank skin compared to black (log2FoldChange = 2.15, adjusted *P*-value = 1.79e-10) or transformed (log2FoldChange = 2.09, adjusted *P*-value = 1.24e-10) skins. Secondly, genes crucial for melanocyte differentiation, such as *WNT16* (Fan et al. 2014), *WNT3A* (Jin et al. 2001; Dunn et al. 2005), migration (*EDNRB2* (Kinoshita et al. 2014)), communication (*MC1R* (Kerje et al. 2003; Schwochow et al. 2021)), melanin production (*TYR* (Chang et al. 2006), *TYRP1*, *DCT* (Li et al. 2019)), packaging (*MLANA* (Hoashi et al. 2005), *PMEL* (Kerje et al. 2004), *SLC45A2* (Gunnarsson et al. 2007)), and transportation (*MLPH* (Vaez et al. 2008)) were either not expressed or significantly less expressed in yellow shank skin, while being highly expressed in black shank skin. Consequently, neither melanoblasts nor melanocytes are present in yellow shank skin, which aligns with the theory of apoptosis affecting melanocyte precursors.

### Mechanisms behind transformed shank formation in chickens

We also incorporated transformed shank samples in our transcriptomic analysis, which revealed differences from black shank samples primarily in genes associated with melanin production, packaging, and transportation, but not in those related to melanocyte differentiation or migration. No indications of apoptosis were observed in these samples. Moreover, the expression levels of *MLANA*, *TYRP1*, *PMEL*, and *MLPH*, while lower than in black shank skin, were still significantly higher in transformed shank skin (transcripts per million (TPM) ranging from 2.2 to 22.5) compared to yellow shank skin. Collectively, these findings suggest that melanocytes are permitted to migrate and differentiate in transformed shank skin, likely due to the *ID*N* allele and reduced *CDKN2A* expression, which mitigates apoptosis. Despite this, visible pigmentation at hatching is absent, possibly due to factors such as *FM*N*. Dorshorst et al. (2011) suggested that *ID*N* facilitates melanoblast migration, while *FM* regulates melanoblast proliferation and maintenance, with the shank potentially being an exception.

### Suitable AAV for in vivo gene editing in chickens

Due to the unique reproductive system of avian species, the application of the powerful gene-editing tool CRISPR/Cas9 in avian species has been less prevalent than in mammals. The first successful CRISPR/Cas9 modification in a chicken was reported in 2016 (Oishi et al. 2016). Consequently, there are fewer resources and less knowledge available regarding suitable AAVs for in vivo gene editing in chickens. We tested three types of AAVs (AAV2, AAV9, AAVdj), but none demonstrated infection of shank dermal cells in chicks. Additionally, the light reflected by shank scales under the microscope adversely affected the observation of fluorescent light. Moving forward, we will test avian-AAV (A3V), as successful transfections in chickens and ducks have been reported (Waldner et al. 2019; Wang et al. 2019).

### Differential expression of melanogenesis genes in cell experiments

Using cell models, we demonstrated that knocking out the candidate sequence or altering *CDKN2A* expression affects melanogenesis pathways and promotes apoptosis, consistent with trends observed in transcriptomic data. Although the upregulation of *CDKN2A* in yellow shank skin (Fig. 3D) is higher than that in our knock-out cells (Fig. 5E), several factors could explain this discrepancy: the low mutation rate (5%) for the double knock-out in black shank primary cells and DF-1 cells, and the fact that DF-1 cells are not fixed for the *ID*N* allele.

Among the well-known melanogenesis genes, only the expression of *TYR* was decreased by knocking out the candidate sequence in *ID*N* cells, and only the expression of *MC1R* was increased by inhibiting *CDKN2A* in *ID*ID* cells. There are two potential reasons for these inconsistencies: 1. Since yellow shank skin lacks melanocyte progenitors, apoptosis may have occurred at earlier stages rather than in the differentiated cells used in our cell experiments. 3. Although apoptosis was observed in our cultured cells, melanogenesis genes could be influenced by other pathways. For example, it has been reported that the chicken Alternate Reading Frame (ARF) protein, encoded by *CDKN2A*, can interact with Mouse Double Minute 2 homolog (MDM2) and protect p53 from degradation (Kim et al. 2003; di Tommaso et al. 2009). Overexpression of p53 in human melanoma cells has been shown to decrease TYR expression and its activity (Kichina et al. 1996). Therefore, increased *CDKN2A* expression could inhibit *TYR* through p53 rather than through a proapoptotic effect in our knocked-out cells.

### Genetic and developmental relationships between ID and B Loci

A remarkable finding from our study is that different mutations of the same gene are responsible for both *ID* and *B*, with these mutations located approximately 0.1 Mb apart, corresponding to a genetic distance of about 0.8 cM (Groenen et al. 2009). The genetic linkage between *ID* and *B* has been extensively studied, as summarized by C. W. Knox (1935), R. C. Punnett (1940), and J. J. Bitgood (1988). According to these earlier studies, the genetic distance between *ID* and *B* varied from 2.9 cM to 13.7 cM. D. C. Warren (1928) argued that the 2.9 cM reported by Serebrovsky and Wassina (1926) might be an underestimation, as the plumage pigment-inhibiting effect of *B*B* could be pleiotropic, potentially inhibiting shank pigmentation and leading to misphenotyping of actual recombinant birds (*B*B*/*ID*N*). Warren (1928) also suggested that the black shank phenotype might result from *E*E* rather than *B*B*, potentially masking the phenotype of other actual recombinant birds (*B*N*/*ID*ID*).

Based on the current understanding of the molecular mechanisms of these mutations, we propose an alternative explanation that the previously reported genetic distances could be overestimations. The *B*B* allele in those linkage studies likely originated from Barred Plymouth Rock, which we now know is the *B1* allele with coding and non-coding SNPs in *CDKN2A*. The ARF protein resulting from this coding mutation has a significantly lower binding affinity to MDM2 than the wild-type (Schwochow Thalmann et al. 2017). Regardless of whether such protein levels are increased by non-coding mutations of *B*B* or *ID*ID*, their low binding affinity might still allow melanocytes to survive, resulting in dark shanks. Consequently, some *B*B*/*ID*ID* birds might have been falsely phenotyped as *B*B*/*ID*N* and counted as crossovers. In other words, *B*B* is expected to promote the expression of dark shanks, contradicting previous observations (Knox 1935; Dorshorst and Ashwell 2009). The previously reported inhibition of shank pigmentation by *B*B* was likely due to *ID*ID*, as they are closely linked (the effect of *B*B*/*ID*ID* varies, with some being dark and others light shanks). Additionally, *B*N*/*ID*N* birds could also be misphenotyped as crossovers due to *FM*N* or *I*I*, which inhibit shank pigmentation (Knox 1935; Zhang et al. 2015).

Furthermore, while the method Serebrovsky and Wassina (1926) used to phenotype shank color is uncertain, it is entirely possible to distinguish epidermal melanin (resulting from *E*E*) from dermal melanin (*ID*N*) through careful observation, as shown in Fig. 1. It is true that epidermal melanin can mask the dermal color, making it difficult to judge the *ID* phenotype. In our mapping population, *E*E* was not involved, leading to more accurate phenotyping of shank color.

In addition to the genetic distance between the *B* and *ID* loci, their cellular functions present an intriguing area of study. Premature differentiation of melanocytes, associated with *CDKN2A*, is implicated in the development of *sex-linked barring*, yet apoptosis was not detected through Caspase-3 assays (Schwochow Thalmann et al. 2017). Conversely, in the current study, Caspase-3 upregulation and apoptosis were observed in light shank chickens and in cells overexpressing *CDKN2A*. The fate of melanocytes, whether they undergo premature differentiation or apoptosis, is likely determined by their developmental stages and the surrounding microenvironment.

Prematurely differentiated melanocytes in feather follicles create periodic patterns on the contour feathers of barring chickens (Schwochow Thalmann et al. 2017). However, the same *B*B* mutation results in a white spot on the head of the chick rather than a periodic pattern (Hellström et al. 2010). Our study demonstrated that, in chicks, the non-periodic pattern on the shank, is associated with the ectopic expression of the same gene. Further research is necessary to confirm the underlying connections.

Schwochow Thalmann *et al*. (2017) demonstrated that the evolution of *sex-linked barring* in chickens is the result of an accumulation of SNPs affecting the expression and function of the same gene. Now, we have added a new functional mutation to this gene, although it is uncertain which mutation occurred first. Therefore, *B* and *ID* alleles provide materials to understand the molecular evolution in a step-by-step manner.

In summary, our findings not only facilitate the molecular breeding of auto-sexing chicken lines but also provide valuable new genetic insights that contribute to a deeper understanding of cellular differentiation, apoptosis, and pattern formation.

## Materials and Methods

### Animals and tissue collections

The animal procedures used in this study were approved by the Institutional Animal Care and Use Committee of Huazhong Agricultural University (HZAUCH-2023-0020).

For observing the inheritance and expression patterns of the shank color, a pure-line Turpan game cock with black shank (*FM*N*/*N*, *ID*N*/*N*) and a White Leghorn hen (*FM*N*/*N*, *ID*ID*/W) with yellow shank from our in-house chicken populations were crossed and 17 offspring were obtained. Their shank colors were recorded and photographed at the age of day 1, 14, and 30, blood samples were collected at day 30. A resource family for *ID* gene mapping was initiated by crossing 1 White Silkie cock (*FM*FM*/*FM*, *ID*N*/*N*) with 1 White Leghorn hen (both are our in-house populations). Then, 3 F_1_ males (*FM*FM*/*N*, *ID*ID*/*N*) and 66 yellow shank Jinghong No.1 commercial layers (*FM*N*/*N*, *ID*ID*/W, purchased from Huadu Yukou Poultry Industry, Beijing, China) were mated and 482 offspring were produced. Shank color of these offspring was recorded at day 1 and week 9, bled at week 9. Blood samples from other 170 Jinghong No.1 commercial layers from the same line were also involved in this study.

In the offspring of the resource family, 8 female day 1 chicks (based on vent sexing) expressing dark shank with similar hatch weight were used for AAV injection experiment. Also in the offspring of the resource family, 12 female chicks at hatch (8 expressing yellow shank and 4 that were dark shank) were euthanized and shank skin tissues were collected for RNA-seq. White Leghorn males and females from our in-house chicken population were also used to produce 60 fertile eggs which were incubated for tissue collection at E12, E14, E16, E18, and at hatch. Dorsal skin tissue from an in-house 60 days old chicken with light shank was collected for cDNA. All above tissue samples were stored in Trizol and at -80℃. Day 15 embryos from Jianghan chicken with dark shank color (n = 16) and White Leghorn (n = 16) from our in-house chicken populations were used for primary cell culture (from shank skin tissues). During tissue collection, these chicks or embryos were also bled for DNA extraction and for genotyping and/or sexing by PCR.

All chicken individuals were raised at the Experimental Poultry Farm at Huazhong Agricultural University and kept in the same conditions.

### DNA extraction and sexing

For each of the blood samples, genomic DNA was extracted by phenol-chloroform method. Other than vent sexing at hatch, sexing was also accomplished using the previously described PCR diagnostic method (Hu et al. 2003).

### Genotyping of the top associated SNP in GWAS

Created Restriction Site-Restriction Fragment Length Polymorphism (CRS-RFLP) was applied to genotype the top associated SNP in GWAS (Supplemental Methods). From the offspring of the resource family, 86 females consisting 64 black shank and 22 yellow shank birds (the phenotypic records were double checked) were selected for genotyping this SNP.

### Scanning for candidate mutations surrounding the top SNP

In the nearly 50 kb region flanking the top SNP (chrZ:78,912,685-78,962,370), 62 pairs of primers (Supplemental Table S5) were designed in the head-to-tail manner. All mutations were investigated in this region by Sanger sequencing of 2 DNA pools. The first pool was constructed by mixing the equal amount of DNA from 10 females of the resource family offspring expressing yellow shank, while the second pool consists 10 black shank females of the offspring (the phenotypic records were double checked).

### RNA extraction and transcriptomic analysis

Total RNA was extracted using Trizol (Invitrogen, Carlsbad, CA, USA) according to the manufacturer’s protocol. The 12 skin samples of day 1 females were extracted for RNA and also genotyped for the top SNP and *FM* mutation (diagnostic test followed “Assay A” and “Assay B” reported previously (Dorshorst et al. 2011)). Thus, the yellow shank individuals at hatch which were wild-type of the top SNP of *ID* and *FM*, were defined as transformed shank (n = 5), others were yellow shank (n = 3) or black shank (n = 4) based on their phenotypes. DEGs between groups were obtained (Supplemental Methods). The expressed genes were defined as genes that TPM > 2 (Wagner et al. 2013). Then KEGG enrichment analyses for the DEGs were done by the clusterProfiler package of R. Venn diagram was generated by the online platform InteractiVenn (http://www.interactivenn.net/) (Heberle et al. 2015).

### Linkage mapping and association studies

Five SNPs located upstream of *CDKN2A* (Supplemental Table S1) were selected since: 1. their allelic differences were large between the above-described 2 DNA pools; 2. they were completely associated with the shank color in the parents (White Silkie cock with dark shank, White Leghorn and Jinghong No.1 commercial layers with light shank). Details of the genotyping methods for these 5 SNPs are listed in Supplemental Table S6. In short, PCR-RFLP was used for genotyping and the samples included 96 yellow shank and 41 black shank females, and 54 yellow shank males in the offspring of the resource family. Females were excluded for analysis if they were heterozygous in at least 1 of the 5 SNPs. Since males could be *ID*ID*/*N* or *ID*ID*/*ID*, only females were used for the linkage mapping, while all males and females were used for association studies.

### Diagnostic test for the candidate SV

The candidate SV was firstly genotyped in the same 86 female individuals used for Sanger sequencing and then in all the 482 offspring of the resource family plus the breed panel (6 breeds, among them, White Leghorn, Jinghong No.1 layers, Houdan are light color on shank, Yunyang Black-bone chickens, Chahua, and Jingyang are dark color on shank. It consists of 288 individuals in total as 48 from each breed, those DNA samples were collected from our previous studies), via PCR and electrophoresis. The primer sequences are, F: 5’-GTGTATTATTCATGGGGTTT-3’, R: 5’-ATCCTGGTACTAAAATTATG-3’, which amplified a 129 bp fragment for a mutant allele or a 272 bp fragment for the reference allele. The genotyping method for the 23 bp insertion was also described (Supplemental Methods).

### WGS data analysis and TF prediction

DNA samples extracted from the blood of 19 Yunyang Da chickens (expressing dark shank, kept in the conservation farm of Yunyang Da chicken in Shiyan City, Hubei Provience, China) were subjected to whole genome sequencing (20X coverage on DNBSEQT7 with 2 × 150 bp paired-end reads). In addition, WGS data from NCBI SRA database, consists of 25 breeds (285 individuals), were also analyzed to genotype the candidate SV (Supplemental Table S2), 13 of them (147 individuals) are known for expressing dark shank color, i.e., red jungle fowl; while 12 of them (138 individuals) are known for expression light shank color, i.e., Leghorn. SNPs and SVs in the candidate SV flanking region were called for all those 304 individuals (Supplemental Methods). Online JASPAR transcription factor database (Rauluseviciute et al. 2024) was applied to predict the putative TF binding sites (Supplemental Methods).

### Reporter assay

Four kinds of recombinant plasmids were constructed, PGL3-basic-reference, PGL3-basic-mutant, PGL3-promoter-reference, and PGL3-promoter-mutant, to investigate the promoter or enhancer activities of the candidate SV. After amplification from genomic DNA, restriction enzyme digestion, cloning into vectors and transfection into DF-1 cells (Supplemental Methods), transcriptional activity was investigated by Dual-Luciferase Reporter Assay System (Promega, Madison, WI, USA) after 48 hours of transcription.

### Expression pattern of CDKN2A

A total of 60 pure line White Leghorn fertile eggs were incubated for tissue collection at E12, E14, E16, E18, and at hatch. As 12 eggs were assigned to each stage, about 10 alive embryo or day 1 chicks for each stage were obtained and then euthanized. After vent or PCR sexing, 3 female embryos for each stage and 4 female chicks at hatch were chosen. For all the embryos and chicks, shank skin tissues were collected, while for the chicks, thigh shin, liver, and leg muscle tissues were also collected. Following total RNA extraction, methods for quantification of mRNA were described (Supplemental Methods). qPCR primer sequences were listed in Supplemental Table S7. Each qPCR assay was carried out using three technical replicates.

### sgRNA-CRISPR/Cas9 system design and construction

From the flanking sequences of the candidate SV (ChrZ:78,942,000-78,943,800), potential target sites were predicted using crispor.tefor.net. Two pairs of target sequences with higher specificity and lower predicted score for off-targets were chosen. For each target sequence, two complementary primers were annealed with the 21-bp target sequence to generate a double-strand DNA with 4-bp overhangs on both ends, and then cloned into the BbsⅠ digested vector pX330-U6-Chimeric_BB-CBh-hSpCas9. Thus, 8 primers for sgRNA annealing were designed (Supplemental Table S8), i.e., 4 types of pX330-U6-sgRNA_BB-CBh-hSpCas9 knock-out vectors were constructed after ligation, transfection, and amplification. Additional details were described (Supplemental Methods).

### Selection for different serotypes of AAV for in vivo injection

We tested 3 serotypes of AAV, AAV-9-EGFP, AAV-DJ-EGFP, and AAV-2-EGFP (Hanbio, Shanghai, China). The 8 female day 1 chicks with similar hatch weight were assigned into 4 groups with 2 chicks in each group. Each chick in Group A, B, and C was injected with AAV-9, AAV-DJ, and AAV-2 respectively, while ddH_2_O was injected into Group D chicks. For each chick, the amount of injection was 10 μL for each time, and each was injected 3 times (1, 7, 14 days). The AAV or ddH_2_O was injected into the dermal of the left shank in the angle of 15°, speed of injection was 1-2 µL/min. The titer for injected AAV was 1×10^12^. The injected chicks were euthanized at day 21 (Supplemental Fig. S3), dermal tissues of the left shank were harvested for cryostat sections. After detecting GFP fluorescent signals, the sections were processed to immunohistochemistry. Additional details were described (Supplemental Methods).

### Construction of double knock-out and CDKN2A knock-down cell lines, and gene expression analysis

Each of the 4 types of vectors was single transfected, or co-transfected into DF-1 cells, for knock-out frequency analysis by electrophoresis. For the best vector combination, amplicons were ligated and transfected into competent cells and knock-out efficiency were analyzed by Sanger sequencing. Then, female (PCR sexing) E15 embryonic shank skin (6 yellow shank and 6 black shank) tissues were collected. Cells were trypsinized, resuspended, strained, seeded, and cultured as primary shank dermal cells. The black shank cells were knocked-out and the yellow shank cells were transfected by siRNA inhibiting chicken *CDKN2A*. Additional details were described (Supplemental Methods). Those modified and original primary cells were extracted for total RNA. Expressions of *CDKN2A* and other pigmentation genes were quantitated. Primer sequences can be found in Supplemental Table S7.

### Overexpressing CDKN2A in cell lines and cell apoptosis assay

The total RNA extracted from the skin of a light shank chicken was reverse-transcripted into cDNA using HiScript II Q RT SuperMix for qPCR (+gDNA wiper) (Vazyme, Nanjing, China). The CDS of *CDKN2A* was amplified from the cDNA, and then inserted into pcDNA3.1 (Genewiz, Suzhou, China). This *CDKN2A* overexpression vector and empty vector pcDNA3.1 were transfected respectively into DF-1 cells via Lipofectamine 3000 (Invitrogen). After 48 hours, these transfected cells were analyzed for the expression of *CDKN2A* and cell apoptosis assay through Annexin V-AbFluor™ 488/PI Apoptosis Detection kit (Abbkine, Wuhan, China). In particular, early apoptotic cells were stained with Annexin V-AbFluor™ 488+/ Propidium Iodide (PI)− and late apoptotic cells were stained with Annexin V-AbFluor™ 488+/ PI+. The stained cells were detected by flow cytometry (Aurora CS, Cytek Biosciences, USA). A total of 10,000 events were recorded for each analysis using FlowJo software.

### Data deposition

The raw RNA sequencing data generated in this study have been submitted to the NGDC database (https://ngdc.cncb.ac.cn/bioproject/) under accession number PRJCA021412. The raw WGS data generated in this study have also been submitted to the NGDC database under accession number PRJCA024727.

## Acknowledgments

We thank Shiyan Kangda Poultry Co. Ltd provided the blood samples of Yunyang Da chickens, and researchers who provided the publicly available WGS data used in this study. We also thank Xuefeng Wang who helped with the construction of the resource family. This work was supported by Major Project of Hubei Hongshan Laboratory (2022hszd006) & Natural Science Foundation of China (No.31772585).

## Author Contributions

S.L. and Q.W. conceived the study. The mapping population was developed by X.Z. and S.L. X.Z. was responsible for phenotyping and conducting GWAS. Q.G. contributed to the linkage mapping and association studies. L.W. was responsible for the bioinformatics analysis. J.C. and H.Y. conducted the flow cytometry assays, while S.Y. was responsible for the remaining functional studies. J.L. conducted the linkage mapping calculations, TF predictions, and WGS data analysis. J.L. also drafted the paper, with contributions from all other authors.

## References

Andersson L, Bed’hom B, Chuong C-M, Inaba M, Okimoto R, Tixier-Boichard M. 2020. The genetic basis for pigmentation phenotypes in poultry. In Advances in Poultry Genetics and Genomics (eds. S.E. Aggrey, H. Zhou, M. Tixier-Boichard, and D.D. Rhoads), pp. 67–106, Burleigh Dodds Science Publishing, Cambridge.

Bateson W, Punnett RC. 1911. The inheritance of the peculiar pigmentation of the silky fowl. J Genet 1: 185–203.

Bitgood JJ. 1988. Linear relationship of the loci for barring, dermal melanin inhibitor, and recessive white skin on the chicken Z chromosome. Poult Sci 67: 530–533.

Cha J, Jin D, Kim JH, Kim SC, Lim JA, Chai HH, Jung S a., Lee JH, Lee SH. 2023. Genome-wide association study revealed the genomic regions associated with skin pigmentation in an Ogye x White Leghorn F2 chicken population. Poult Sci 102: 102720.

Chakraborty S, Utter MB, Frias MA, Foster DA. 2021. Cancer cells with defective RB and CDKN2A are resistant to the apoptotic effects of rapamycin. Cancer Lett 522: 164–170.

Chang CM, Coville JL, Coquerelle G, Gourichon D, Oulmouden A, Tixier-Boichard M. 2006. Complete association between a retroviral insertion in the tyrosinase gene and the recessive white mutation in chickens. BMC Genomics 7: 1–15.

di Tommaso A, Hagen J, Tompkins V, Muniz V, Dudakovic A, Kitzis A, Ladeveze V, Quelle DE. 2009. Residues in the alternative reading frame tumor suppressor that influence its stability and p53-independent activities. Exp Cell Res 315: 1326–1335.

Dickinson ME, Flenniken AM, Ji X, Teboul L, Wong MD, White JK, Meehan TF, Weninger WJ, Westerberg H, Adissu H, et al. 2016. High-throughput discovery of novel developmental phenotypes. Nature 537: 508–514.

Dorshorst B, Molin AM, Rubin CJ, Johansson AM, Strömstedt L, Pham MH, Chen CF, Hallböök F, Ashwell C, Andersson L. 2011. A complex genomic rearrangement involving the Endothelin 3 locus causes dermal hyperpigmentation in the chicken. PLoS Genet 7: e1002412.

Dorshorst B, Okimoto R, Ashwell C. 2010. Genomic regions associated with dermal hyperpigmentation, polydactyly and other morphological traits in the Silkie chicken. J Hered 101: 339–350.

Dorshorst BJ, Ashwell CM. 2009. Genetic mapping of the sex-linked barring gene in the chicken. Poult Sci 88: 1811–1817.

Dunn KJ, Brady M, Ochsenbauer-Jambor C, Snyder S, Incao A, Pavan WJ. 2005. WNT1 and WNT3a promote expansion of melanocytes through distinct modes of action. Pigment Cell Res 18: 167–180.

Eriksson J, Larson G, Gunnarsson U, Bed’hom B, Tixier-Boichard M, Strömstedt L, Wright D, Jungerius A, Vereijken A, Randi E, et al. 2008. Identification of the Yellow skin gene reveals a hybrid origin of the domestic chicken. PLoS Genet 4: e1000010.

Fan YP, Wang P, Fu WX, Dong T, Qi C, Liu L, Guo G, Li C, Cui XG, Zhang SL, et al. 2014. Genome-wide association study for pigmentation traits in Chinese Holstein population. Anim Genet 45: 740–744.

Foissac S, Djebali S, Munyard K, Vialaneix N, Rau A, Muret K, Esquerré D, Zytnicki M, Derrien T, Bardou P, et al. 2019. Multi-species annotation of transcriptome and chromatin structure in domesticated animals. BMC Biol 17: 108.

Groenen MA, Wahlberg P, Foglio M, Cheng HH, Megens HJ, Crooijmans RPMA, Besnier F, Lathrop M, Muir WM, Wong GKS, et al. 2009. A high-density SNP-based linkage map of the chicken genome reveals sequence features correlated with recombination rate. Genome Res 19: 510–519.

Gunnarsson U, Hellström AR, Tixier-Boichard M, Minvielle F, Bed’hom B, Ito S, Jensen P, Rattink A, Vereijken A, Andersson L. 2007. Mutations in SLC45A2 cause plumage color variation in chicken and Japanese quail. Genetics 175: 867–877.

Heberle H, Meirelles VG, da Silva FR, Telles GP, Minghim R. 2015. InteractiVenn: A web-based tool for the analysis of sets through Venn diagrams. BMC Bioinformatics 16: 1–7.

Hellström AR, Sundström E, Gunnarsson U, Bed’Hom B, Tixier-Boichard M, Honaker CF, Sahlqvist AS, Jensen P, Kämpe O, Siegel PB, et al. 2010. Sex-linked barring in chickens is controlled by the CDKN2A/B tumour suppressor locus. Pigment Cell Melanoma Res 23: 521–530.

Hoashi T, Watabe H, Muller J, Yamaguchi Y, Vieira WD, Hearing VJ. 2005. MART-1 is required for the function of the melanosomal matrix protein PMEL17/GP100 and the maturation of melanosomes. J Biol Chem 280: 14006–14016.

Hu RY, Geng X, Ma J, Chen YS, Li ZK, Ding XY. 2003. A simple and universal method for molecular sexing of birds. Shi Yan Sheng Wu Xue Bao 36: 401–404.

Ibarrola-Villava M, Fernandez LP, Pita G, Bravo J, Floristan U, Sendagorta E, Feito M, Avilés JA, Martin-Gonzalez M, Lázaro P, et al. 2010. Genetic analysis of three important genes in pigmentation and melanoma susceptibility: CDKN2A, MC1R and HERC2/OCA2. Exp Dermatol 19: 836–844.

Ji ZG, Huo CY, Yang PQ. 2020. Genistein inhibited the proliferation of kidney cancer cells via CDKN2a hypomethylation: role of abnormal apoptosis. Int Urol Nephrol 52: 1049–1055.

Jin EJ, Erickson CA, Takada S, Burrus LW. 2001. Wnt and BMP signaling govern lineage segregation of melanocytes in the avian embryo. Dev Biol 233: 22–37.

Kang XT, Song SF, Wang Y Bin, Huang YQ, Xiong YH, Zhu XJ, Yang YL. 2002. Study on shank color, plumage color and feather growth of slow-feathering pure line of Gushi fowl. J Northwest Sci-tech Univ Agric For Sci Ed 30: 57–59.

Kerje S, Lind J, Schütz K, Jensen P, Andersson L. 2003. Melanocortin 1-receptor (MC1R) mutations are associated with plumage colour in chicken. Anim Genet 34: 241–248.

Kerje S, Sharma P, Gunnarsson U, Kim H, Bagchi S, Fredriksson R, Schütz K, Jensen P, Von Heijne G, Okimoto R, et al. 2004. The Dominant white, Dun and Smoky color variants in chicken are associated with insertion/deletion polymorphisms in the PMEL17 gene. *Genetics* **168**: 1507–1518.

Kichina J, Green A, Rauth S. 1996. Tumor suppressor p53 down-regulates tissue-specific expression of tyrosinase gene in human melanoma cell lines. Pigment Cell Res 9: 85–91.

Kim SH, Mitchell M, Fujii H, Llanos S, Peters G. 2003. Absence of p16INK4a and truncation of ARF tumor suppressors in chickens. Proc Natl Acad Sci 100: 211–216.

Kinoshita K, Akiyama T, Mizutani M, Shinomiya A, Ishikawa A, Younis HH, Tsudzuki M, Namikawa T, Matsuda Y. 2014. Endothelin receptor B2 (EDNRB2) is responsible for the tyrosinase-independent recessive white (mow) and mottled (mo) plumage phenotypes in the chicken ed. A. Roulin. PLoS One 9: e86361.

Knox CW. 1935. The inheritance of shank color in chickens. Genetics 20: 529–544.

Li GQ, Li DF, Yang N, Qu LJ, Hou ZC, Zheng JX, Xu GY, Chen SR. 2014. A genome-wide association study identifies novel single nucleotide polymorphisms associated with dermal shank pigmentation in chickens. Poult Sci 93: 2983–2987.

Li JY, Bed’hom B, Marthey S, Valade M, Dureux A, Moroldo M, Péchoux C, Coville JL, Gourichon D, Vieaud A, et al. 2019. A missense mutation in TYRP1 causes the chocolate plumage color in chicken and alters melanosome structure. Pigment Cell Melanoma Res 32: 381–390.

Lizio M, Deviatiiarov R, Nagai H, Galan L, Arner E, Itoh M, Lassmann T, Kasukawa T, Hasegawa A, Ros MA, et al. 2017. Systematic analysis of transcription start sites in avian development ed. J. Briscoe. PLOS Biol 15: e2002887.

Luan Y, Zhang W, Xie J, Mao J. 2021. CDKN2A inhibits cell proliferation and invasion in cervical cancer through LDHA-mediated AKT/mTOR pathway. Clin Transl Oncol 23: 222– 228.

Makino S, Fukaya T. 2024. Dynamic modulation of enhancer-promoter and promoter-promoter connectivity in gene regulation. BioEssays 1–7.

Oishi I, Yoshii K, Miyahara D, Kagami H, Tagami T. 2016. Targeted mutagenesis in chicken using CRISPR/Cas9 system. Sci Rep 6: 23980.

Perini F, Cendron F, Lasagna E, Cassandro M, Penasa M. 2024. Genomic insights into shank and eggshell color in Italian local chickens. Poult Sci 103: 103677.

Punnett RC. 1940. Genetic studies in poultry - XI. The legbar. J Genet 41: 1–8.

Rauluseviciute I, Riudavets-Puig R, Blanc-Mathieu R, Castro-Mondragon JA, Ferenc K, Kumar V, Lemma RB, Lucas J, Chèneby J, Baranasic D, et al. 2024. JASPAR 2024: 20th anniversary of the open-access database of transcription factor binding profiles. Nucleic Acids Res 52: D174–D182.

Sanyal A, Lajoie BR, Jain G, Dekker J. 2012. The long-range interaction landscape of gene promoters. Nature 489: 109–113.

Schwochow D, Bornelöv S, Jiang TX, Li JY, Gourichon D, Bed’Hom B, Dorshorst B, Chuong CM, Tixier-Boichard M, Andersson L. 2021. The feather pattern autosomal barring in chicken is strongly associated with segregation at the MC1R locus. Pigment Cell Melanoma Res 34: 1015–1028.

Schwochow Thalmann D, Ring H, Sundström E, Cao X, Larsson M, Kerje S, Höglund A, Fogelholm J, Wright D, Jemth P, et al. 2017. The evolution of Sex-linked barring alleles in chickens involves both regulatory and coding changes in CDKN2A. PLoS Genet 13: e1006665.

Serebrovsku AS, Wassina ET. 1926. On the topography of the sex-chromosome in fowls. J Genet 17: 211–216.

Smyth JR. 1990. Genetics of plumage, skin and eye pigmentation in chickens. In Poultry Breeding and Genetics (ed. R.D. Crawford), Elsevier, Amsterdam.

Szabo Q, Bantignies F, Cavalli G. 2019. Principles of genome folding into topologically associating domains. Sci Adv 5: eaaw1668.

Tian M, Hao R, Fang SY, Wang YQ, Gu XR, Feng CG, Hu XX, Li N. 2014. Genomic regions associated with the sex-linked inhibitor of dermal melanin in Silkie chicken. Front Agric Sci Eng 1: 242–249.

Vaez M, Follett SA, Bed’hom B, Gourichon D, Tixier-Boichard M, Burke T. 2008. A single point-mutation within the melanophilin gene causes the lavender plumage colour dilution phenotype in the chicken. BMC Genet 9: 1–9.

Wagner GP, Kin K, Lynch VJ. 2013. A model based criterion for gene expression calls using RNA-seq data. Theory Biosci 132: 159–164.

Waldner DM, Visser F, Fischer AJ, Bech-Hansen NT, Stell WK. 2019. Avian Adeno-associated viral transduction of the postembryonic chicken retina. Transl Vis Sci Technol 8: 1–13.

Wang AP, Liu L, Gu LL, Wu S, Guo CM, Feng Q, Xia WL, Yuan C, Zhu SY. 2019. Expression of duck hepatitis A virus type 1 VP3 protein mediated by avian adeno-associated virus and its immunogenicity in ducklings. Acta Virol 63: 53–59.

Warren DC. 1928. Sex-linked characters of poultry. Genetics 13: 421–433.

Wu GQ, Wang X, Zhou HY, Chai KQ, Xue Q, Zheng AH, Zhu XM, Xiao JY, Ying XH, Wang FW, et al. 2015. Evidence for transcriptional interference in a dual-luciferase reporter system. Sci Rep 5: 1–10.

Xu JG, Lin SD, Gao XF, Nie QH, Luo Q Bin, Zhang XQ. 2017. Mapping of Id locus for dermal shank melanin in a Chinese indigenous chicken breed. J Genet 96: 977–983.

Yang N, Jiang RS. 2005. Recent advances in breeding for quality chickens. Worlds Poult Sci J 61: 373–382.

Yin HG, Zeng ZJ, Pan YM, Zhou KY. 2001. The genetic expression of dark shank in chicken. Sichuan Anim Vet Sci 28: 17–18.

Zhang XD, Wang HH, Zhang CX, Li QH, Chen XH, Lou LF. 2015. Analysis of skin color change and related gene expression after crossing of Dongxiang black chicken and ISA layer. Genet Mol Res 14: 11551–11561.

Zhu F, Yin ZT, Zhao QS, Sun YX, Jie YC, Smith J, Yang YZ, Burt DW, Hincke M, Zhang ZD, et al. 2023. A chromosome-level genome assembly for the Silkie chicken resolves complete sequences for key chicken metabolic, reproductive, and immunity genes. Commun Biol 6: 1–15.

